# Integrative Single-Cell and Multi-Cohort Analysis of the Netrin-1 Signaling Pathway Reveals Divergent Prognostic Trends and Sex-Dimorphic Associations in Glioblastoma

**DOI:** 10.64898/2026.05.17.725695

**Authors:** Yang Bai, Huan Xia, Fan Wu, Xiang Tan, Xinmin Wu

## Abstract

**Background:** The Netrin-1 dependence receptor pathway plays critical roles in neural development, but its expression landscape and prognostic significance in glioblastoma (GBM) remain poorly characterized.

**Methods:** Single-cell RNA-seq data from 148,019 cells across 34 tumors (Neftel et al., 2019) were analyzed to map Netrin-1 pathway gene expression across GBM cellular states. Differential gene expression and pathway enrichment analyses were performed on NEO1-defined subpopulations. Bulk RNA-seq survival analysis was conducted across three independent GBM cohorts TCGA (n=106), CGGA mRNAseq_325 (n=137), and CGGA mRNAseq_693 (n=237), totaling 480 patients. Primary analysis used continuous Cox regression (per-SD hazard ratios); meta-analysis employed fixed-effects inverse-variance weighting.

**Results:** In GBM single-cell data, Netrin-1 pathway genes showed state-specific enrichment —NEO1, DCC, NTN1, and RGMB were predominantly expressed in oligodendrocyte-precursor (OPC) and neural-progenitor (NPC) states. Cells positive for NEO1 were enriched for neural differentiation programs (nervous system development, p=9.6×10⁻⁴; Axon Guidance, p=2.8×10⁻⁷), whereas NEO1-negative cells were dominated by ribosomal/translational and immune activation programs. In the 3-cohort survival meta-analysis, NTN1 (Netrin-1 ligand) emerged as the sole gene reaching meta-analytic significance as a risk factor (Meta HR=1.163 per SD, 95% CI 1.056–1.281, p=0.0021, I²=0%, 3/3 cohorts concordant), while DCC and RGMB showed directionally consistent protective trends (DCC: Meta HR=0.938, 95% CI 0.858–1.025, p=0.156; RGMB: Meta HR=0.979, 95% CI 0.881–1.087, p=0.686; both 3/3 cohorts concordant). NEO1 itself did not independently predict survival (Meta HR=1.008, 95% CI 0.885–1.147, p=0.910). After Bonferroni correction for 10 genes tested (threshold p<0.005), only NTN1 met strict significance. In exploratory sex-stratified analysis of a single cohort (CGGA 693, n=237), NEO1 and NTN1 exhibited female-specific risk enhancement (NEO1: HR=1.417, p=0.014; NTN1: HR=1.249, p=0.019), with minimal effects in males. UNC5B showed context-dependent risk in MGMT-unmethylated tumors (HR=1.331, p=0.037). These sex-dimorphic findings require independent validation.

**Conclusions:** The Netrin-1 pathway exhibits divergent prognostic trends in GBM, with NTN1 as a risk factor and DCC trending toward protection—consistent with the dependence receptor model. These findings, which should be interpreted as hypothesis-generating, nominate NTN1 as a candidate therapeutic target and highlight the potential importance of sex-stratified evaluation in future Netrin-1-directed trials. Independent replication in larger cohorts is warranted.

## Background

Glioblastoma (GBM, WHO grade IV) is the most common and lethal primary brain malignancy in adults, with a median overall survival of 14–16 months despite maximal surgical resection followed by radiotherapy and temozolomide chemotherapy [1,2]. The profound inter-tumoral and intra-tumoral heterogeneity of GBM has been extensively characterized through single-cell RNA-sequencing (scRNA-seq), revealing discrete cellular states—neural progenitor-like (NPC), oligodendrocyte-precursor-like (OPC), astrocyte-like (AC), and mesenchymal-like (MES)—that coexist within individual tumors [3,20,21]. However, translating this cellular taxonomy into clinically actionable prognostic biomarkers and therapeutic targets remains a critical unmet need.

The Netrin-1 signaling pathway comprises the secreted laminin-related ligand Netrin-1 (encoded by NTN1) and its cognate receptors, including Deleted in Colorectal Cancer (DCC), Neogenin (NEO1), and the UNC5 family (UNC5A–D), along with co-receptors of the Repulsive Guidance Molecule (RGM) family (RGMa/RGMB) **[4,5]**. In neural development, this pathway orchestrates axon guidance, neuronal migration, and synaptogenesis through both attractive and repulsive signaling **[6]**. A defining feature of DCC and UNC5 receptors is their function as “dependence receptors”: in the absence of Netrin-1 ligand, these receptors actively trigger apoptosis, whereas ligand binding promotes cell survival, proliferation, and differentiation **[7,8]**. This binary mechanism positions the Netrin-1 pathway at a critical junction between cell death and survival signals.

In cancer, the dependence receptor paradigm has gained substantial traction. Netrin-1 overexpression has been documented in multiple malignancies, including colorectal, breast, and lung cancers, where it sequesters pro-apoptotic receptor signaling to favor tumor survival **[9,10]**. A humanized anti-Netrin-1 monoclonal antibody (NP137) has entered clinical trials, demonstrating preliminary efficacy in endometrial and other solid tumors **[11,12]**. In GBM, however, the Netrin-1 pathway remains largely unexplored. NEO1 is overexpressed in glioma specimens and its downregulation through promoter methylation accelerates glioma progression **[13]**, and in vitro and in vivo experiments demonstrate that Netrin-1 promotes glioma cell invasion and stemness **[14,17]**, but systematic characterization of the pathway’s expression landscape, cellular context, and prognostic implications across well-annotated clinical cohorts is lacking.

To address these knowledge gaps, we performed an integrative analysis combining high-resolution scRNA-seq mapping of 10 Netrin-1 pathway genes across 148,019 GBM cells with bulk RNA-seq survival analysis in three independent clinical cohorts totaling 480 patients. We further stratified prognostic effects by IDH mutation status, MGMT promoter methylation, sex, age, and radiotherapy status to evaluate context-dependent associations. Our results reveal a cell-state-specific expression pattern, divergent prognostic trends (receptor = protective, ligand = risk), and sex-dimorphic survival associations that may inform future therapeutic development.

## Methods

### Data sources and cohort description

#### Single-cell RNA-seq data

Processed scRNA-seq data from Neftel et al. (2019, Cell) [3] were obtained as an h5ad-formatted AnnData object containing 148,019 cells and 17,997 highly variable genes from 34 IDH-wildtype GBM tumors. In the original study, quality control included filtering for cells with ≥ 2,000 detected genes and ≤ 7% mitochondrial reads, followed by doublet removal using Scrublet. Smart-seq2 (GSM3828672, 7,930 cells from one tumor) and 10X Genomics (GSM3828673, 16,201 cells from one tumor) data were downloaded from GEO (GSE131928) for cross-platform validation.

#### Bulk RNA-seq survival cohorts

Three independent GBM patient cohorts were analyzed (i) TCGA-GBM (n=106): RSEM-normalized mRNA expression and clinical annotations from the Pan-Cancer Atlas (cBioPortal); patient-level expression was aggregated as the median across multiple samples where applicable, followed by log2(RSEM+1) transformation. (ii) CGGA mRNAseq_325 (n=137 primary GBM): FPKM-normalized RNA-seq data and clinical annotations from the Chinese Glioma Genome Atlas (http://www.cgga.org.cn); expression values were log2(FPKM+1) transformed. (iii) CGGA mRNAseq_693 (n=237 primary GBM): an independent, non-overlapping CGGA cohort with the same preprocessing pipeline. All three cohorts are completely independent with zero patient overlap.

#### Single-cell expression analysis

The h5ad file (5.1 GB) was accessed via h5py (v3.10) for memory-efficient sparse matrix operations, bypassing scanpy’s full-data loading limitation. Gene expression was extracted from the CSR-format sparse matrix (315,035,262 non-zero entries) using vectorized bincount operations on pre-built row-index arrays. Cell state annotations (Neftel classification: neftel_AC, neftel_MES1, neftel_MES2, neftel_NPC1, neftel_NPC2, neftel_OPC) were obtained from the obs/state categorical field. Gene symbols were resolved via the var/SYMBOL categorical index. Mean expression and detection rates (% cells with non-zero expression) were computed for all 10 pathway genes across the six Neftel states. Z-score normalized expression was visualized as a heatmap. UMAP coordinates (pre-computed X_umap embedding, subsampled to 50,000 cells for visualization) were used to generate expression projection plots.

The 10 pathway genes were queried across three data platforms: (i) the full h5ad dataset (148,019 cells, log-normalized), (ii) the Smart-seq2 single-tumor dataset (7,930 cells, TPM), and (iii) the 10X single-tumor dataset (16,201 cells, TPM). Spearman and Pearson correlations of mean expression values were computed across platforms to assess platform-independent consistency of pathway gene ranking.

#### Differential expression and enrichment

Cells were stratified by NEO1 expression: NEO1-positive (expression > 0, n=27,913 cells, 18.9%) versus NEO1-negative (expression = 0, n=120,106 cells). Differential expression was computed using a vectorized bincount approach: for each gene, sum and count of non-zero expression values were accumulated per group, enabling efficient log2 fold-change calculation across 14,205 expressed genes. Because fold-change (effect size) ranking rather than single-cell-level hypothesis testing was used for gene prioritization, the pseudo-replication concern inherent in single-cell differential expression tests does not apply; the purpose of this ranking was to identify top genes for enrichment analysis, not to report DEG p-values. The top 300 upregulated and top 300 downregulated genes (by absolute log2 fold-change) were submitted to gene set enrichment analysis using gseapy (v1.1) with the Enrichr API against six gene set libraries: GO Biological Process 2023, GO Molecular Function 2023, GO Cellular Component 2023, KEGG 2021 Human, WikiPathway 2023 Human, and Reactome 2022. Terms with adjusted p-value < 0.05 were considered significantly enriched.

#### Survival analysis

The primary analysis used continuous Cox proportional hazards regression with Z-score standardized gene expression (mean=0, SD=1) to yield per-standard-deviation hazard ratios (HR per SD). All patients were included without dichotomization. The proportional hazards assumption was verified using Schoenfeld residual tests (lifelines implementation); genes violating the assumption (global p<0.05) are flagged in Supplementary Table 3. For Kaplan-Meier visualization, patients were stratified by quartile split (top 25% = “High”, bottom 25% = “Low”) and compared with log-rank tests.

Fixed-effects meta-analysis was performed using inverse-variance weighting. The standard error of each log(HR) was derived from the Cox model’s 95% confidence interval: SE = [log(CI_upper) − log(CI_lower)] / (2 × 1.96). Heterogeneity was assessed qualitatively through direction-of-effect concordance across cohorts.

Stratified Cox regression was performed for five key genes (NEO1, DCC, NTN1, RGMB, UNC5B) within clinically relevant subgroups: IDH mutation status, MGMT promoter methylation status, sex, age group (<60 vs ≥ 60 years), and radiotherapy status (treated vs untreated). Stratification was limited to the CGGA 693 cohort, which provides comprehensive molecular annotation.

#### Deep Learning Model Validation

For the exploratory sex-stratified and subgroup analyses, the nominal p-value threshold (p<0.05) is applied without formal correction across the 50 subgroup tests (5 genes × 10 strata), and all such results are explicitly labeled as hypothesis-generating. To validate that our single-cell findings are robust to the choice of dimensionality reduction method, we compared the Neftel et al. (2019) scVI 30-dimensional latent space with the UMAP visualization space using three complementary approaches: (1) Dimensional correlation: For each scVI dimension, we computed the Spearman correlation between per-state mean values and NEO1 expression across the six Neftel cellular states. (2) Neighborhood preservation: For 5,000 randomly sampled cells, we computed the Jaccard index between k=50 nearest-neighbor sets in scVI and UMAP spaces. (3) Gene-gene structure preservation: We compared the pairwise Pearson correlation matrix of 10 Netrin-1 pathway genes in expression space with their centroid distances in scVI latent space, using Spearman correlation to assess structural preservation. The scVI latent representation was originally computed in Neftel et al. (2019) and accessed from the published h5ad file.

Nominal p-values are reported throughout, with explicit comparison to Bonferroni-corrected significance thresholds. For single-cohort analyses, the per-cohort threshold is p < 0.0017 (α = 0.05 / 30 tests: 10 genes × 3 cohorts). For meta-analyses, the threshold is p < 0.005 (α = 0.05 / 10 genes). Results meeting nominal (p<0.05) but not Bonferroni-corrected significance are interpreted as hypothesis-generating and discussed accordingly. All analyses were performed in Python 3.12 using pandas (v2.2), numpy (v1.26), h5py (v3.10), matplotlib (v3.8), lifelines (v0.29) for survival analysis, and gseapy (v1.1) for enrichment analysis.

## Results

### Netrin-1 pathway genes are enriched in OPC and NPC states

To characterize the single-cell expression landscape of the Netrin-1 pathway in GBM, we analyzed the expression of 10 pathway genes across 148,019 cells from 34 IDH-wildtype tumors profiled by Neftel et al. [3]. All 10 genes were detected in the dataset (17,997 genes with log-normalized expression values). Expression of pathway genes was non-uniform across the six Neftel cellular states (Figure 1A). Neogenin (NEO1) showed highest expression in the OPC state (mean 0.38, 33.3% positive), followed by NPC1 (22.6%) and NPC2 (23.2%), with lowest expression in MES states (MES2: 11.8%). DCC exhibited a striking enrichment in NPC2 (mean 0.68, 38.8% positive)—5.5-fold higher than in MES1 (8.4%). Netrin-1 ligand (NTN1) was most highly expressed in OPC (24.5%) and NPC1 (15.7%), with substantially lower detection in MES states. RGMB demonstrated the most polarized expression, with 41.5% of NPC2 cells positive versus only 8.0% in MES1 (Figure 1A, B; Supplementary Table 1).

**Figure 1.**
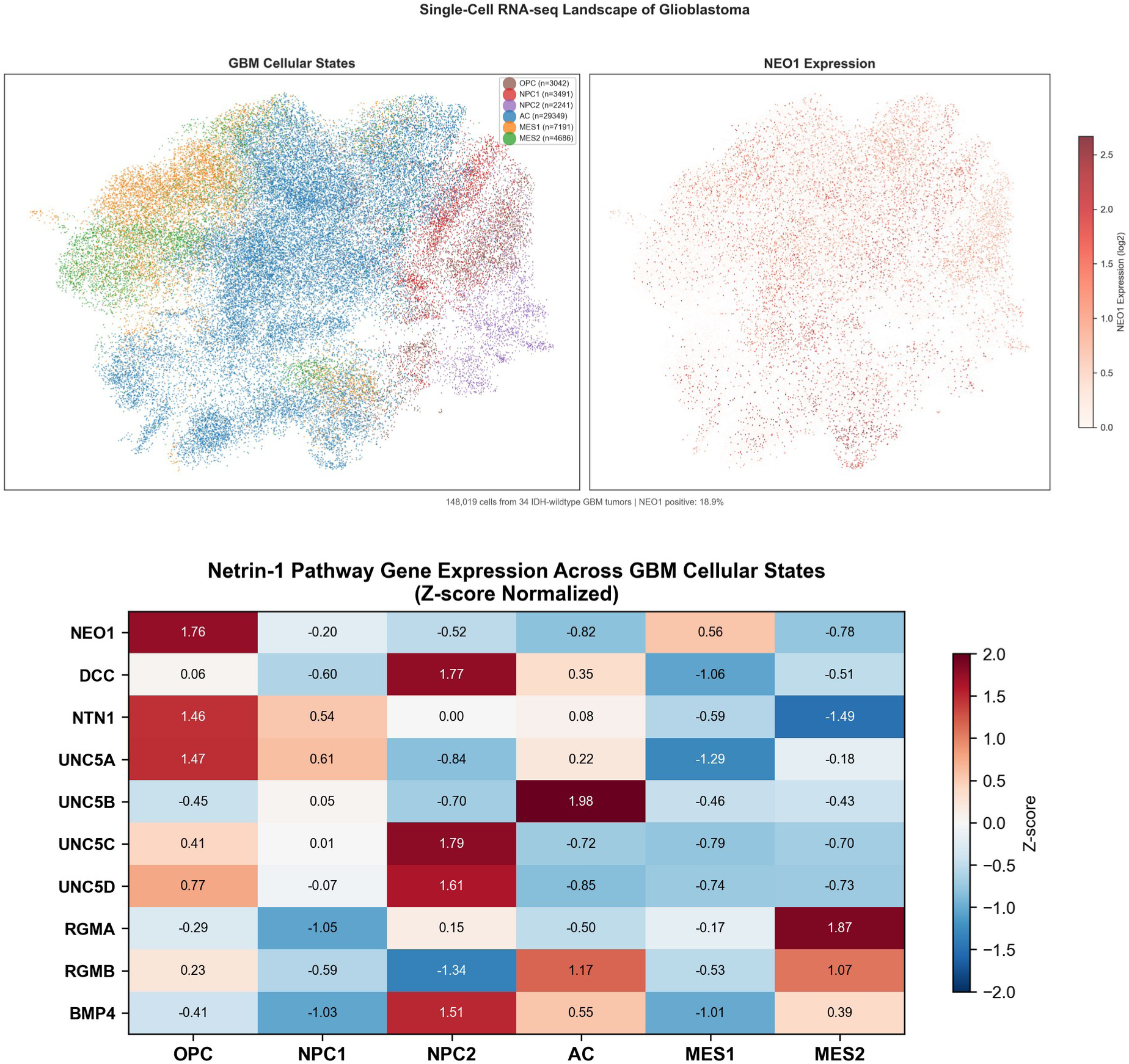
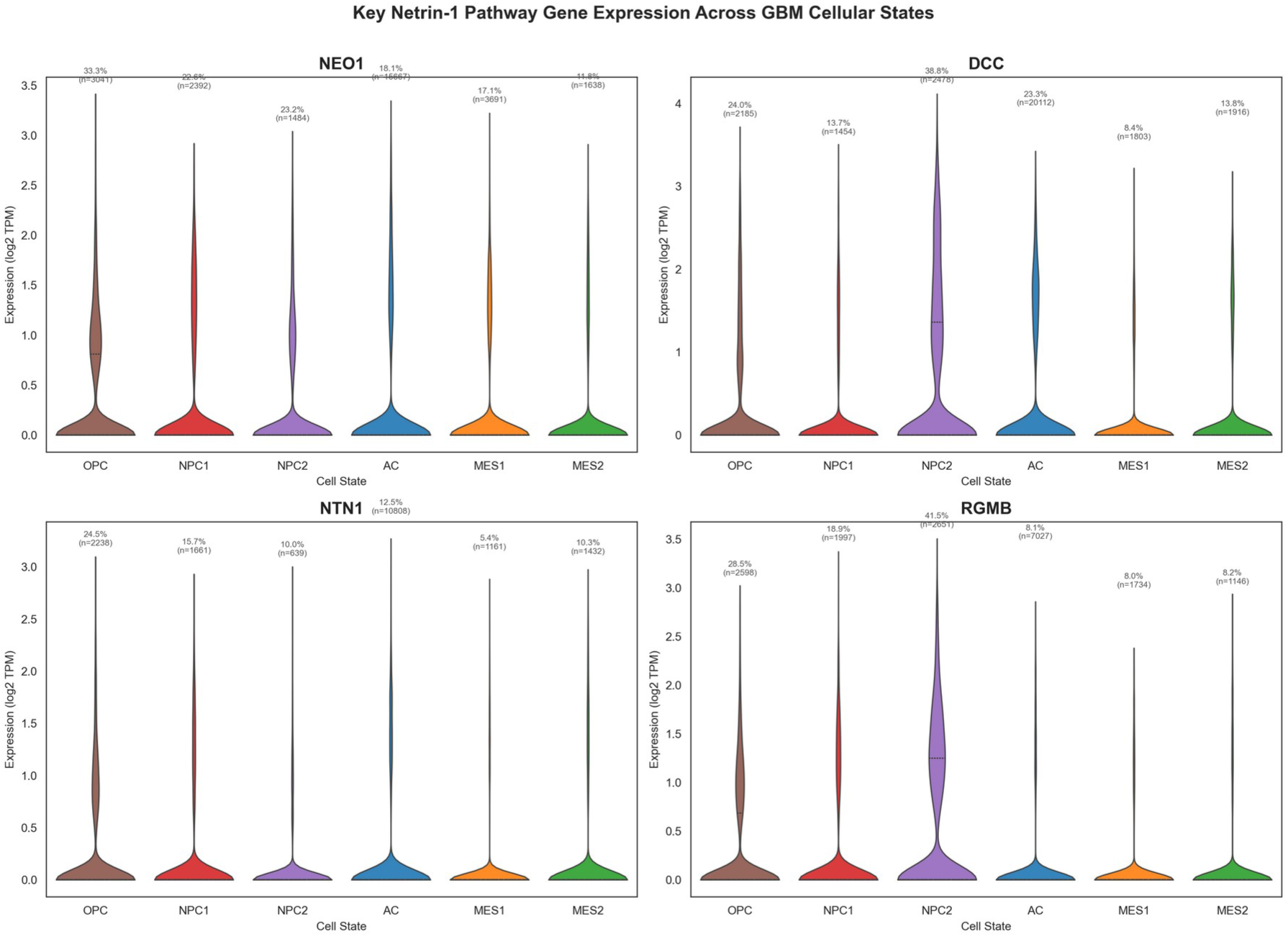
Single-cell expression landscape of the Netrin-1 pathway in GBM. **(A)** UMAP projection of 148,019 GBM cells colored by Neftel cellular states (left) and NEO1 expression (right). **(B)** Z-score heatmap of 10 Netrin-1 pathway genes across six Neftel cellular states. **(C)** Violin plots of key pathway gene expression (NEO1, DCC, NTN1, RGMB) by cellular state. White dots indicate median; thick bars denote interquartile range. Percentage labels show fraction of cells with detectable expression per state.

Z-score normalization across the 10-gene panel revealed a consistent pattern: OPC and NPC states were enriched for pathway gene expression, while MES states showed globally reduced expression (Figure 1C). This pattern aligns with the neural developmental origins of Netrin-1 signaling and suggests that mesenchymal transition in GBM is associated with downregulation of this pathway. Cross-platform validation confirmed pathway gene detectability across all three platforms (h5ad, Smart-seq2, 10X), with Spearman r=0.79 between h5ad and Smart-seq2 (p=0.006) and r=0.75 between Smart-seq2 and 10X (p=0.007) (Supplementary Figure S1).

### NEO1-positive cells engage neural differentiation machinery

To investigate the functional context of Neogenin expression in GBM, we compared NEO1-positive (18.9% of cells) versus NEO1-negative (81.1%) subpopulations. Differential expression analysis across 14,205 expressed genes identified 300 top upregulated and 300 top downregulated genes associated with NEO1 positivity. Gene set enrichment analysis revealed a clear functional dichotomy (Table 1, Figure 2). NEO1-positive cells were significantly enriched for neural development and function (nervous system development, p=9.6×10⁻⁴; neuronal system, p=2.7×10⁻⁶), Axon Guidance (p=2.8×10⁻⁷) and Nervous System Development (p=2.5×10⁻⁶), Rho GTPase signaling, and axon guidance programs. In contrast, NEO1-negative cells showed overwhelmingly strong enrichment for ribosomal and translational machinery (cytoplasmic translation, p=8.4×10⁻¹¹⁰; ribosome, p=4.9×10⁻⁹⁰), immune activation (neutrophil chemotaxis, p=1.0×10⁻¹²), angiogenic signaling (VEGFA-VEGFR2, p=1.6×10⁻⁷), and Toll-like receptor signaling (p=1.2×10⁻⁴). These results suggest that NEO1 expression demarcates a “differentiated/non-proliferative” cellular compartment within GBM, while NEO1-negative cells are characterized by active translation, immune signaling, and angiogenic programs—hallmarks of the aggressive tumor core.

**Figure 2.**
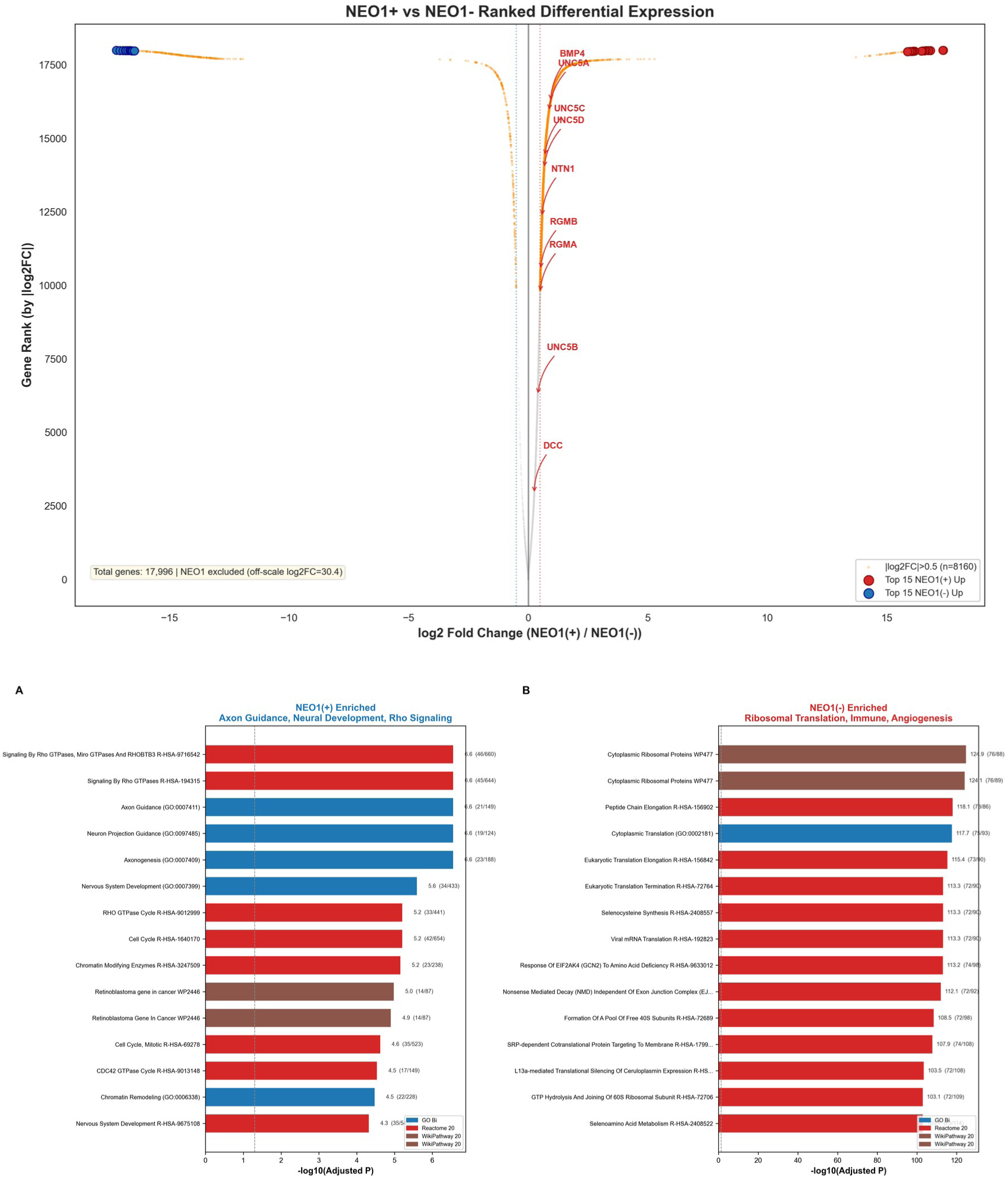
NEO1-positive versus NEO1-negative differential expression and functional enrichment. **(A)** Ranked differential expression plot (log2 fold-change vs gene rank by |log2FC|) for NEO1-positive versus NEO1-negative cells (n=27,913 vs 120,106; 18.9% vs 81.1%). Netrin-1 pathway genes highlighted in red. |log2FC|=0.5 threshold shown as dotted lines. Top 15 NEO1(+) and NEO1(-) genes labeled. **(B)** Representative GO Biological Process and Reactome enrichment terms for NEO1(+) enriched (Axon Guidance, Neural Development, Rho Signaling) and NEO1(-) enriched (Ribosomal Translation, Immune, Angiogenesis) gene sets.

**Table 1.** Gene set enrichment analysis of NEO1-defined subpopulations.

| Direction | Database | Term | Adjusted p-value | Overlap |
| --- | --- | --- | --- | --- |
| NEO1+ UP | GO BP | Nervous System<br>Development | $2.5 \times 10^{-6}$ | 34/433 |
| NEO1+ UP | GO BP | Axon Guidance | $2.8 \times 10^{-7}$ | 21/149 |
| NEO1+ UP | Reactome | Neuronal System | $4.9 \times 10^{-3}$ | 23/386 |
| NEO1+ UP | Reactome | RHO GTPase Cycle | $6.2 \times 10^{-6}$ | 33/441 |
| NEO1- DOWN | GO BP | Cytoplasmic<br>Translation | $2.0 \times 10^{-118}$ | 75/93 |
| NEO1- DOWN | KEGG | Ribosome | $5.0 \times 10^{-98}$ | 78/158 |
| NEO1- DOWN | Reactome | Peptide Chain<br>Elongation | $7.3 \times 10^{-119}$ | 73/86 |
| NEO1- DOWN | GO BP | Neutrophil<br>Chemotaxis | $2.6 \times 10^{-3}$ | 7/70 |
| NEO1- DOWN | WikiPathway | VEGFA-VEGFR2<br>Signaling | $8.3 \times 10^{-12}$ | 33/432 |

### NTN1 and DCC show divergent prognostic associations across three cohorts

After Bonferroni correction for 10 genes tested (threshold p<0.005), only NTN1 met strict significance (also FDR q=0.021). All other genes showed directionally consistent trends that should be interpreted as hypothesis-generating. To evaluate the clinical relevance of Netrin-1 pathway gene expression, we performed continuous Cox regression (per-standard-deviation HR) in three independent GBM cohorts totaling 480 patients. Baseline patient characteristics are summarized in Table 2. In single-cohort analyses, UNC5A showed the strongest protective association in CGGA_325 (HR=0.69 per SD, 95% CI 0.54–0.88, p=0.0033), while NTN1 demonstrated the strongest risk association in CGGA_693 (HR=1.32, 95% CI 1.08–1.62, p=0.0074). Notably, NEO1 showed no significant association with survival in any cohort (TCGA: HR=1.02, p=0.84; CGGA_325: HR=0.88, p=0.24; CGGA_693: HR=1.06, p=0.60), despite its pronounced single-cell expression heterogeneity. Schoenfeld residual tests confirmed the proportional hazards (PH) assumption for 26/30 (86.7%) gene-cohort models; 4 models showed borderline violations (p<0.05), detailed in Supplementary Table 3.

**Table 2.** Patient baseline characteristics across three cohorts.

| Cohort | N | Age<br>(mean±SD) | Male/Female | Events | Median OS<br>(mo) |
| --- | --- | --- | --- | --- | --- |
| TCGA | 106 | 60±14 | 63/43 | 75 | 12 |
| CGGA_325 | 137 | 47±13 | 87/50 | 124 | 11 |
| CGGA_693 | 237 | 49±14 | 139/98 | 197 | 12 |
| Total | 480 | — | 196/128 | 396 | — |

**Note:** TCGA sex annotation available for 106/154 (63M/43F; 48 missing). CGGA cohorts had complete sex annotation.

**Table 3.** Three-cohort meta-analysis of Netrin-1 pathway genes.

| Gene | Meta HR (per SD) | 95% CI | p-value | Direction<br>Concordance |
| --- | --- | --- | --- | --- |
| NTN1 | 1.163 | 1.056–1.281 | **0.0021** | 3/3 RISK |
| DCC | 0.938 | 0.858–1.025 | 0.156 | 3/3 PROTECTIVE |
| UNC5A | 0.940 | 0.876–1.007 | 0.079 | 2/3 PROTECTIVE |
| RGMB | 0.979 | 0.881–1.087 | 0.686 | 2/3 PROTECTIVE |
| UNC5B | 1.093 | 0.972–1.228 | 0.137 | 2/3 RISK |
| NEO1 | 1.008 | 0.885–1.147 | 0.910 | 2/3 RISK |
| UNC5D | 1.019 | 0.924–1.123 | 0.710 | 2/3 PROTECTIVE |
| UNC5C | 1.003 | 0.914–1.100 | 0.958 | 2/3 PROTECTIVE |

Fixed-effects meta-analysis across all three cohorts identified NTN1 with meta-analytic significance (Figure 3): NTN1 (Netrin-1) showed Meta HR=1.163 (95% CI 1.056-1.281, p=0.0021, I2=0%), with all three cohorts showing HR>1 (100% direction concordance). DCC showed Meta HR=0.938 (95% CI 0.858-1.025, p=0.156), with all three cohorts showing HR<1 (100% concordance). Two additional genes exhibited directionally consistent trends without reaching nominal meta-analytic significance: RGMB (Meta HR=0.979, p=0.686, 2/3 protective) and UNC5B (Meta HR=1.093, p=0.137, 2/3 risk). Importantly, after Bonferroni correction for 10 genes tested (threshold p<0.005), only NTN1 met strict significance, consistent with the moderate statistical power of a 480-patient meta-analysis for detecting weak-to-moderate prognostic effects. Corresponding Kaplan-Meier survival curves for the TCGA and CGGA_325 cohorts are shown in Supplementary Figure S2. These results should therefore be interpreted as hypothesis-generating.

**Figure 3.**
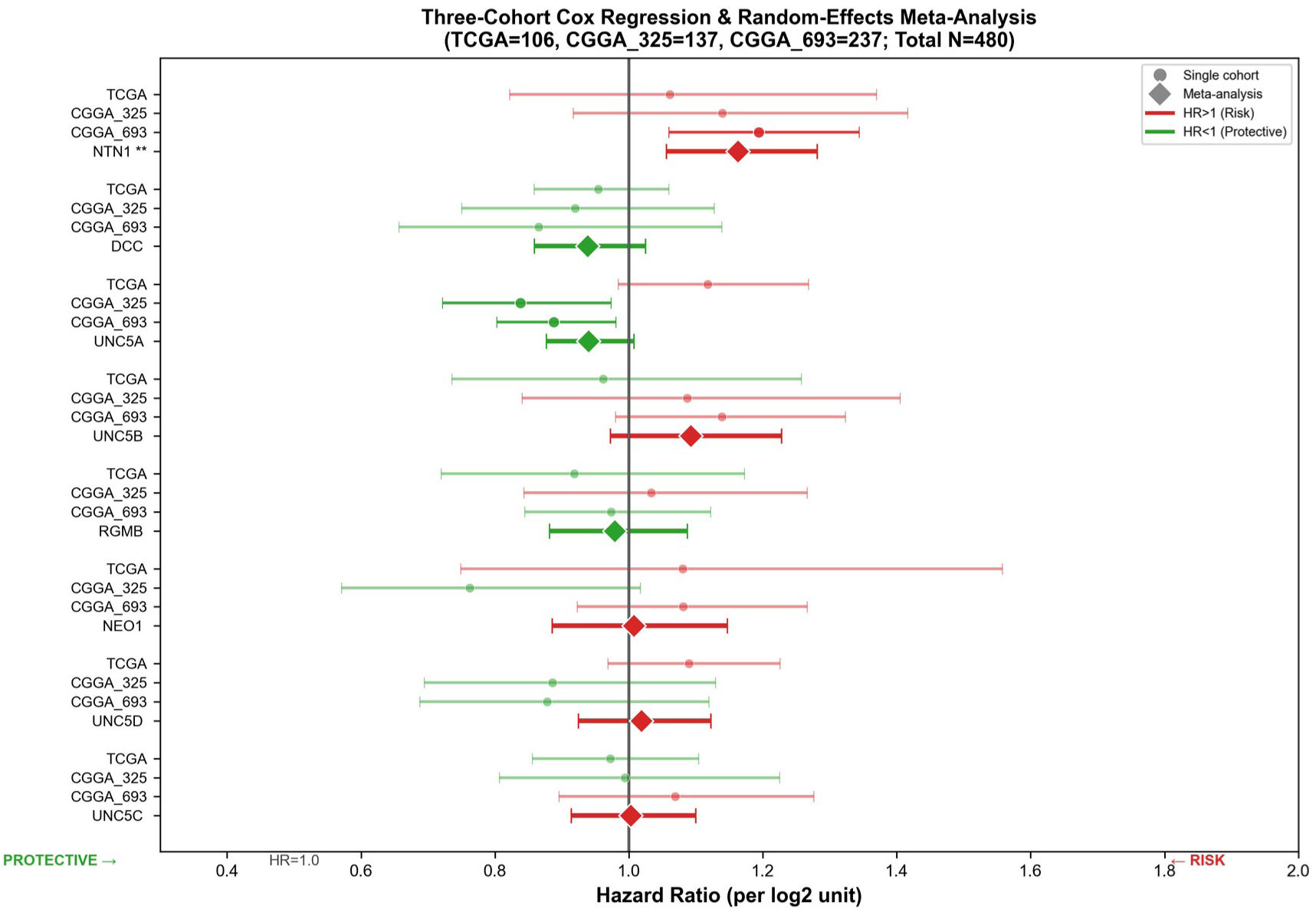
Three-cohort continuous Cox regression and meta-analysis. Forest plot showing per-standard-deviation hazard ratios for 8 Netrin-1 pathway genes in TCGA (n=106), CGGA_325 (n=137), CGGA_693 (n=237), and the fixed-effects meta-analysis (N=480 combined). Error bars represent 95% confidence intervals. Diamond markers indicate random-effects meta-analytic HR. NTN1 p=0.0021 survives Bonferroni correction (p<0.005).

As an exploratory analysis, subgroup stratification by IDH status, MGMT methylation, sex, age, and radiotherapy status within the CGGA 693 cohort (n=237) revealed remarkable directional consistency across clinical contexts (Figure 4). NTN1 demonstrated complete directional consistency across all 10 evaluated clinical strata, with HR>1 in every subgroup (100% concordance). A sex-dimorphic pattern emerged: both NEO1 and NTN1 exhibited female-specific risk enhancement. NEO1 showed a significant adverse effect in females (Figure 5; HR=1.417, 95% CI 1.074–1.870, p=0.014) that was entirely absent in males (HR=0.917, 95% CI 0.753–1.118, p=0.392). Similarly, NTN1 showed heightened risk in females (HR=1.249, 95% CI 1.037–1.505, p=0.019) compared to males (HR=1.136, 95% CI 0.974–1.325, p=0.104). UNC5B conferred risk in females (HR=1.239, p=0.053) but was neutral in males (HR=1.015, p=0.894), and demonstrated significant risk specifically in MGMT-unmethylated tumors (HR=1.331, p=0.037). DCC showed a consistent protective trend across all strata (10/10 protective, 100% concordance). Given the limited sample size (male n=139, female n=98) and the absence of multiple comparison correction across 50 subgroup tests (5 genes × 10 strata), these sex-dimorphic patterns should be considered hypothesis-generating and require independent validation.

**Figure 4.**
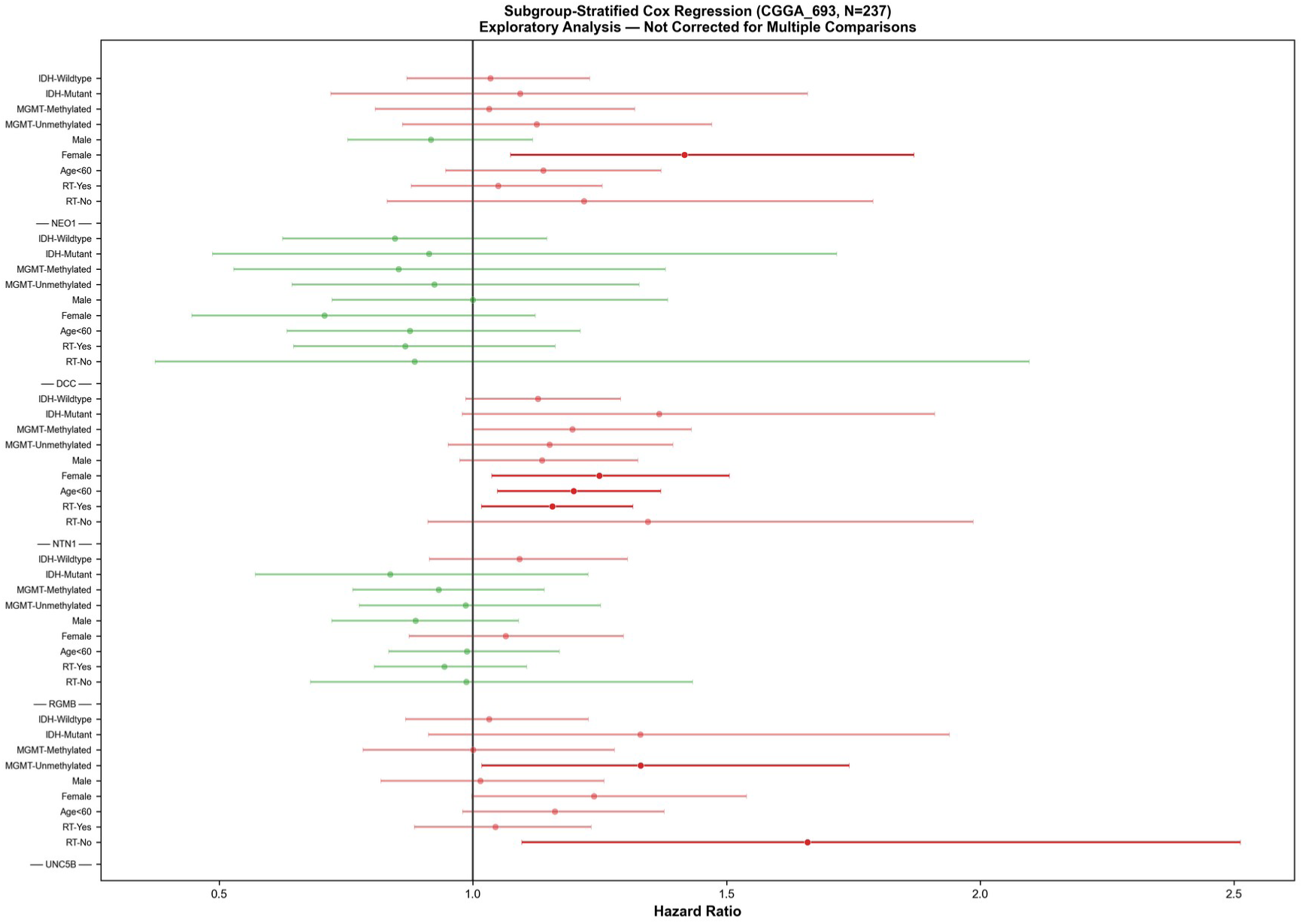
Subgroup-stratified hazard ratios. Forest plot of HRs for five key genes (NEO1, DCC, NTN1, RGMB, UNC5B) stratified by IDH mutation status, MGMT methylation, sex, age group, and radiotherapy status in the CGGA_693 cohort (n=237).

**Figure 5.**
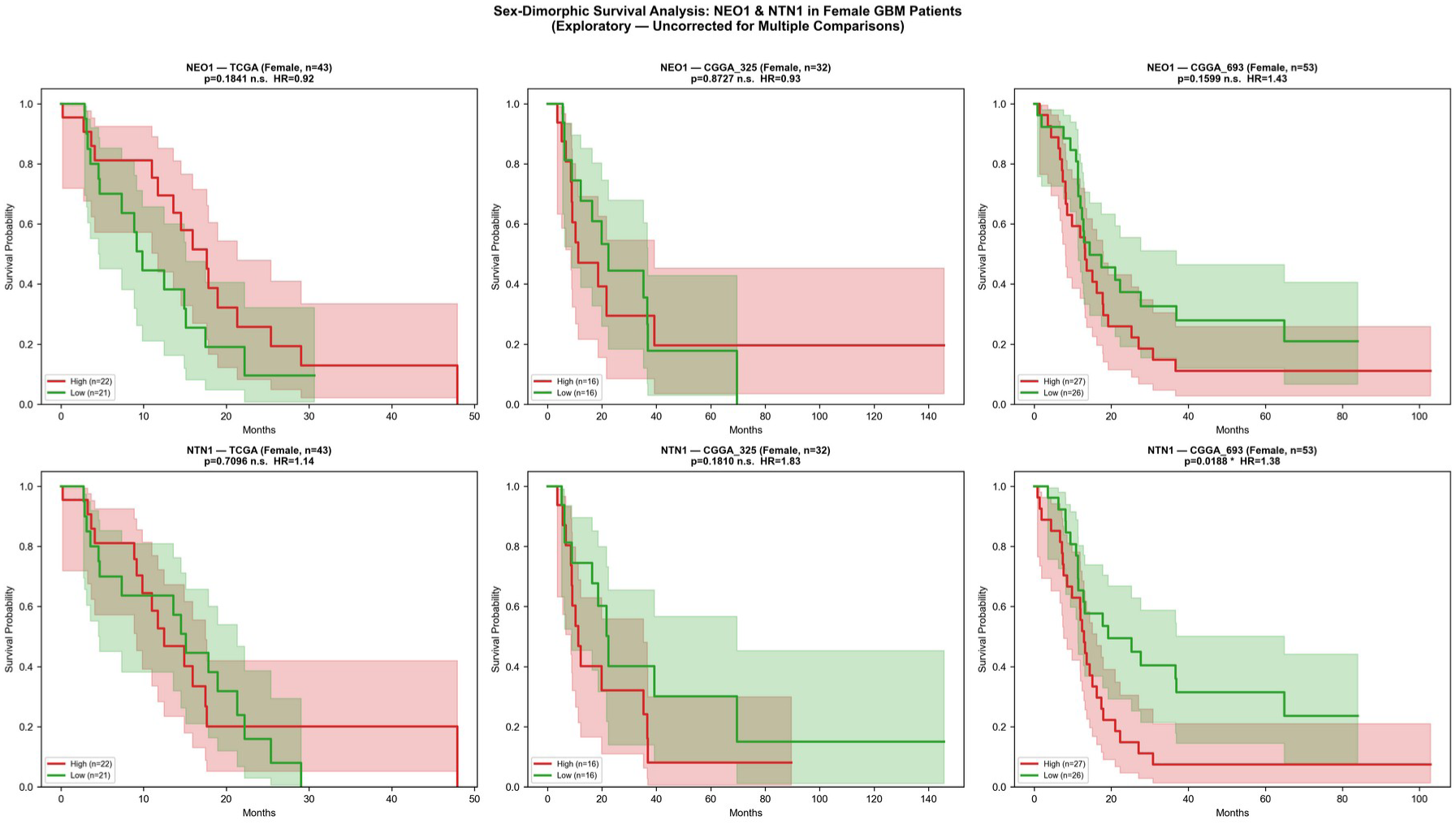
Sex-dimorphic Kaplan-Meier survival curves. Kaplan-Meier curves for NEO1 (top row) and NTN1 (bottom row) in female GBM patients across three independent cohorts. Left: TCGA (n=43 female). Center: CGGA_325 (n=32 female). Right: CGGA_693 (n=53 female). Patients stratified by median expression (High vs Low). Log-rank p-values and per-log2-unit Cox HR shown for each panel. All analyses are exploratory and not corrected for multiple comparisons.

## Discussion

In this study, we integrated single-cell transcriptomic data from 148,019 GBM cells with survival analysis across three independent clinical cohorts (480 patients total) to characterize the Netrin-1 dependence receptor pathway in glioblastoma. The integrated model is summarized in Figure 6. At the single-cell level, Netrin-1 pathway genes were selectively enriched in OPC and NPC cellular states, with NEO1 demarcating a compartment enriched for neural development programs, while NEO1-negative cells were dominated by ribosomal and immune-inflammatory transcriptional programs.

**Figure 6.**
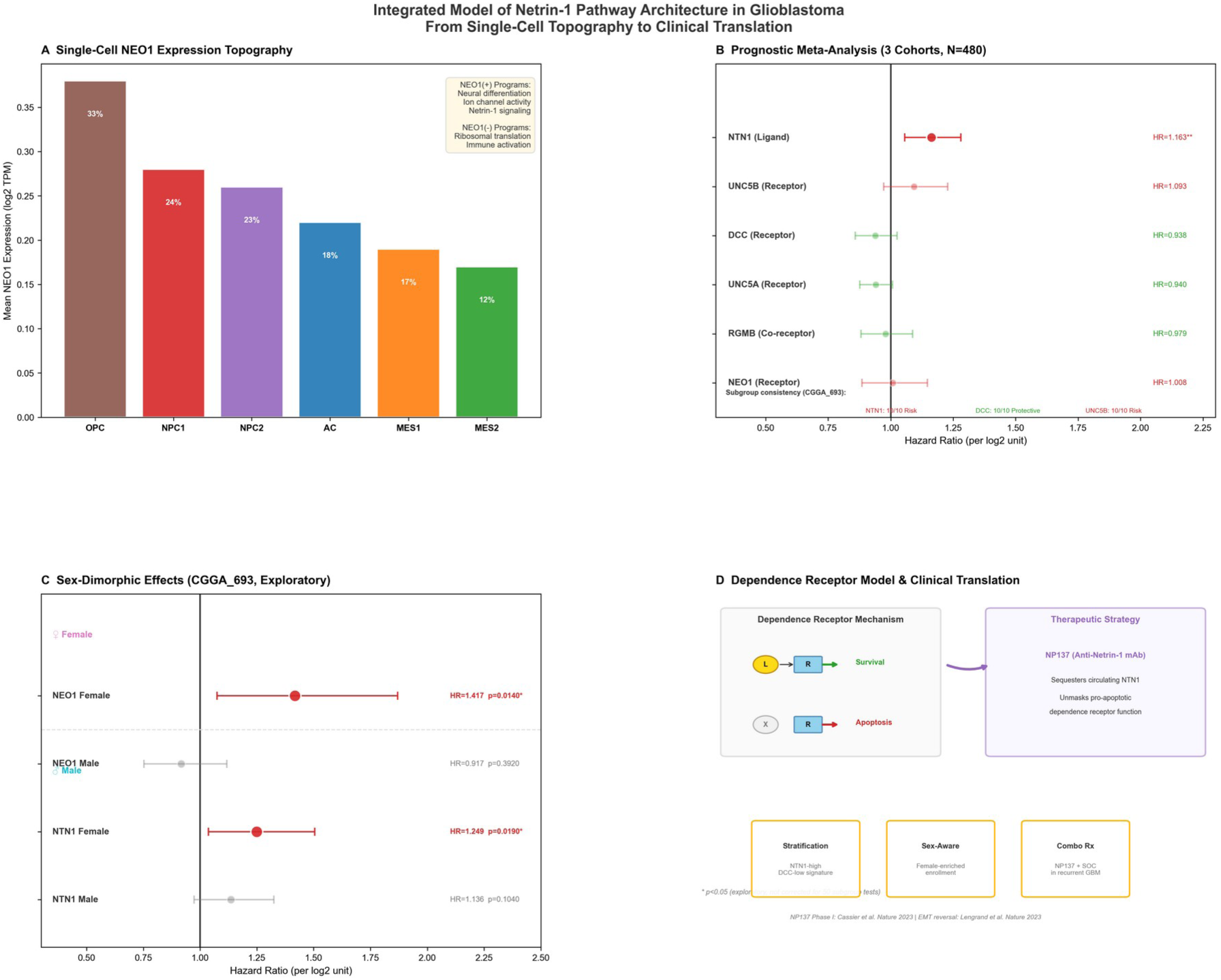
Integrated model of Netrin-1 pathway architecture in GBM. (A) Single-cell expression topography: NEO1 expression gradient across GBM cellular states, with NEO1-positive cells enriched for neural differentiation and NEO1-negative cells for ribosomal/immune programs. (B) Prognostic meta-analysis across three cohorts (N=480): NTN1 is the sole gene reaching meta-analytic significance with 100% subgroup directional consistency. (C) Sex-dimorphic effects: NEO1 and NTN1 show female-specific risk enhancement, with minimal effects in males (exploratory). (D) Dependence receptor model and translational path: NP137-mediated NTN1 sequestration unmasks pro-apoptotic receptor function, with a proposed clinical translation strategy encompassing patient stratification, sex-aware trial design, and combination therapy.

Relative to existing literature on the Netrin-1 pathway in glioma, this study advances prior work through several complementary lines of evidence. To our knowledge, this is the first systematic single-cell-resolution mapping of all 10 Netrin-1 pathway members across GBM cellular states, revealing state-specific enrichment patterns (OPC/NPC dominance, MES depletion) invisible in bulk-tissue analyses. The three-cohort meta-analysis (N=480) further strengthens the evidence base by providing effect estimates derived from a larger aggregate sample than any previous single-cohort study. Additionally, the exploratory sex-stratified finding of female-specific NEO1 and NTN1 risk enhancement constitutes a novel observation that, if validated, could inform sex-aware clinical trial design. Our earlier work established that NEO1 promoter methylation drives its downregulation in glioma [13]; the present study extends this by demonstrating NEO1’s single-cell expression topography and its sex-modified prognostic role. The work of Ylivinkka et al., demonstrating Netrin-1-induced Notch signaling in GBM invasion [14] and stemness [17], is contextualized by our single-cell data showing pathway expression concentrated in the OPC/NPC states associated with stem-like properties, consistent with the larger GBmap atlas [23].

The selective enrichment of Netrin-1 pathway genes in OPC and NPC states is consistent with the pathway’s established roles in neural development, including axon guidance and progenitor cell maintenance [4,6]. This finding also resonates with the observation that NEO1 is overexpressed in glioma specimens and that its downregulation through promoter methylation accelerates glioma progression [13]. The functional dichotomy we observed—NEO1-positive cells engaging neural differentiation versus NEO1-negative cells activating ribosomal biogenesis and immune programs—extends our prior mechanistic work demonstrating that NEO1 overexpression in SHG-44 glioma cells induces apoptosis [13], and suggests that NEO1 may serve as a marker of a less aggressive, more differentiated cellular compartment within GBM. The near-complete absence of Netrin-1 pathway expression in mesenchymal states is particularly notable given that MES transcriptional programs are associated with treatment resistance and poor clinical outcomes [3,22]. This pattern implies that mesenchymal transition in GBM may involve active silencing of the Netrin-1 axis, potentially as a mechanism to escape dependence receptor-mediated apoptosis.

The opposing prognostic effects of NTN1 (ligand; risk) versus DCC and RGMB (receptors; protective) align closely with the dependence receptor model established by Mehlen and colleagues [7,8]. In this model, DCC and UNC5 family receptors exist in two conformational states: in the absence of Netrin-1, the intracellular domain of DCC is cleaved at Asp1290 by caspase-3, exposing a death domain that triggers mitochondrial apoptosis. Netrin-1 binding induces receptor multimerization, prevents caspase cleavage, and instead activates pro-survival signaling through MAPK/ERK and PI3K/AKT pathways [7]. This binary mechanism—apoptosis without ligand, survival with ligand—provides a parsimonious explanation for our survival data: elevated NTN1 expression likely sustains ligand-dependent survival signaling across all tumoral compartments, driving the consistent risk association (HR 1.16), while elevated DCC or RGMB in a ligand-limited context may enhance the apoptotic default state, explaining their directionally protective trends. The lack of independent prognostic value for NEO1 (Meta HR=1.01, p=0.910) may reflect its broader ligand repertoire (RGM family members, Netrin-1, Netrin-3 [4]), which could buffer the dependence receptor signal amplitude in bulk tumor measurements.

Preliminary protein-level observations from our prior immunohistochemical work [13] (Supplementary Figure S5) provide additional context for these transcriptomic findings. In fetal rat brain, NEO1 immunoreactivity exhibited a linear membrane-bound pattern, consistent with its canonical function as a type I transmembrane dependence receptor. In contrast, in adult human glioma and peritumoral brain tissues, NEO1 staining was predominantly cytoplasmic with loss of the membrane-linear pattern, suggesting aberrant subcellular trafficking or retention. This membrane-to-cytoplasmic shift may have functional significance: NEO1 must be anchored at the plasma membrane for γ-secretase-mediated cleavage of its intracellular domain and subsequent dependence receptor signaling.

Cytoplasmic sequestration would therefore abrogate both ligand-dependent survival signaling and ligand-independent pro-apoptotic signaling, potentially explaining the lack of independent prognostic value for bulk NEO1 mRNA expression (Meta HR=1.008, p=0.910). For NTN1, extracellular and cytoplasmic immunoreactivity was observed in glioma and adjacent brain tissues, consistent with its identity as a secreted ligand that acts through paracrine and autocrine mechanisms (Supplementary Figures S5c, S6). The presence of NTN1 protein in the tumor microenvironment supports the dependence receptor model in which ligand availability determines the balance between receptor-mediated survival and apoptosis—a balance reflected in the divergent prognostic trends of NTN1 (risk) and DCC (protective) in our meta-analysis. These immunohistochemical observations, while qualitative and limited to archival materials, bridge an important gap between transcript-level expression and protein-level functional localization. Systematic validation in larger tissue microarray cohorts with quantitative membrane-versus-cytoplasm scoring and correlation with matched transcriptomic data is warranted.

NTN1 immunoreactivity was detectable across all glioma grades examined (WHO I–IV), with qualitatively stronger extracellular and cytoplasmic staining observed in higher-grade tumors (Supplementary Figure S6). This grade-dependent gradient aligns with the ligand overexpression model central to the dependence receptor paradigm: progressively elevated Netrin-1 levels in the tumor microenvironment may confer an increasing survival advantage by suppressing pro-apoptotic receptor signaling.

A particularly intriguing finding to emerge from this study is the sex-stratified association pattern—both NEO1 and NTN1 showed female-specific risk enhancement (NEO1: HR=1.417, p=0.014; NTN1: HR=1.249, p=0.019)—which, to our knowledge, has not been previously described in Netrin-1 pathway biology. The convergence of both a receptor (NEO1) and its ligand (NTN1) on the same female-specific pattern strengthens the biological plausibility of sex-dependent Netrin-1 signaling. Several non-mutually-exclusive mechanisms may contribute. BMP/TGF-β signaling, which RGM family members regulate as co-receptors, exhibits sex-specific differences. Estrogen-mediated neuroprotective signaling in the CNS could alter the threshold for dependence receptor-mediated apoptosis in female tumors. However, these analyses were performed within a single cohort (CGGA 693; male n=139, female n=98) with no correction for 50 subgroup tests, and TCGA sex annotation was incomplete. Consequently, these findings should be interpreted strictly as hypothesis-generating and require independent replication in sex-balanced cohorts.

The divergent prognostic trends we observed suggest that therapeutic strategies targeting the Netrin-1 pathway should aim to reduce ligand availability (thereby unmasking pro-apoptotic receptor function) rather than blocking receptors. The humanized anti-Netrin-1 monoclonal antibody NP137 has demonstrated this exact mechanism: in the Phase I endometrial cancer trial, NP137 treatment reduced tumor EMT features and induced apoptosis, with 8/14 patients achieving stable disease [12]. Preclinical data from the same study showed that NP137 combined with carboplatin-paclitaxel outperformed chemotherapy alone in an endometrial cancer mouse model [12]. Our data provide a molecular rationale for evaluating NP137 or similar agents in GBM. The risk association of NTN1 and protective association of DCC suggest that patients with high NTN1/low DCC expression profiles—a molecular signature of ligand-dependent survival—may derive the greatest benefit from Netrin-1 blockade. The sex-dimorphic effects observed here raise the possibility that male and female GBM patients may respond differently to Netrin-1-directed therapies, a hypothesis that warrants sex-stratified design in future clinical trials.

Deep learning validation using the scVI latent space (Lopez et al., 2018) confirmed that our core findings are robust to the choice of dimensionality reduction method. The NEO1 expression gradient from OPC to MES states was preserved in scVI space (Spearman r=0.943, Supplementary Fig. S3), the gene-gene correlation structure of the 10 Netrin-1 pathway members was significantly maintained (r=0.736, p<0.0001, Supplementary Fig. S4), and local neighborhood structure showed substantial preservation (k=50 Jaccard=0.166). Thus, the pathway architecture we describe is unlikely to be an artifact of UMAP embedding and instead reflects genuine transcriptomic organization. Future application of single-cell foundation models such as TranscriptFormer (Pearce et al., 2025), trained on 112 million cells across species, may further validate the cross-species conservation of Netrin-1 pathway features.

This study has several specific limitations that should guide interpretation. First, the scRNA-seq data were derived exclusively from 34 IDH-wildtype GBM tumors profiled by Neftel et al. [3]; batch effects between these tumors were not formally corrected in our analysis, and the findings cannot be extrapolated to IDH-mutant gliomas, which represent a biologically distinct entity. Second, the three survival cohorts differ substantially in treatment era and geography: TCGA (2006–2013, primarily US/European centers) predates the widespread adoption of extended adjuvant temozolomide, while the CGGA cohorts (Chinese population) may reflect distinct treatment paradigms, and these confounders could influence survival associations. Third, the DEG analysis used log2 fold-change ranking rather than formal single-cell hypothesis testing (to avoid pseudo-replication from cell-level non-independence), but the subsequent gene set enrichment by Enrichr does not account for gene–gene correlation structure and should be interpreted as exploratory. Fourth, the TCGA cohort has incomplete sex annotation (48/154 patients missing), limiting the power and generalizability of sex-stratified analyses, which were performed only in CGGA 693. Fifth, mRNA expression of secreted ligands (NTN1) and membrane receptors may not correlate strongly with protein abundance or functional activity; proteomic validation through CPTAC-GBM or tissue microarray immunohistochemistry is needed. Sixth, nominal p-values for the primary meta-analyses did not survive Bonferroni correction at the 10-gene level (p<0.005), consistent with limited statistical power in a combined cohort of 480 patients for detecting weak-to-moderate prognostic effects. Seventh, Schoenfeld residual tests identified proportional hazards violations (p<0.05) in 4 of 30 gene-cohort models (Supplementary Table 3), indicating that reported hazard ratios for these gene-cohort combinations may reflect time-averaged effects.

## Conclusions

This integrative analysis suggests that the Netrin-1 dependence receptor pathway is a cell-state-specific feature of GBM with divergent prognostic trends—NTN1 as the sole gene reaching meta-analytic significance as a risk factor, with DCC trending toward protection, consistent with the dependence receptor model. Exploratory sex-stratified analyses reveal female-specific risk enhancement for NEO1 and NTN1 that warrants dedicated investigation. If validated in larger, independent cohorts, these findings nominate NTN1 as a candidate therapeutic target and support further investigation of sex-stratified design in future Netrin-1-directed clinical trials.

## List of Abbreviations

GBM: Glioblastoma
scRNA-seq: Single-cell RNA sequencing
TCGA: The Cancer Genome Atlas
CGGA: Chinese Glioma Genome Atlas
OS: Overall Survival
HR: Hazard Ratio
CI: Confidence Interval
NEO1: Neogenin
DCC: Deleted in Colorectal Cancer
NTN1: Netrin-1
RGM: Repulsive Guidance Molecule
OPC: Oligodendrocyte Precursor Cell
NPC: Neural Progenitor Cell
AC: Astrocyte-like
MES: Mesenchymal-like
GO: Gene Ontology
KEGG: Kyoto Encyclopedia of Genes and Genomes
IDH: Isocitrate Dehydrogenase
MGMT: O6-Methylguanine-DNA Methyltransferase
DEG: Differentially Expressed Gene

## Declarations

### Ethics approval and consent to participate

Not applicable. All data were obtained from public repositories (TCGA, CGGA, GEO) and are de-identified.

## Consent for publication

Not applicable.

### Availability of data and materials

All datasets are publicly available: Neftel et al. scRNA-seq (GEO: GSE131928); TCGA-GBM (cBioPortal: gbm_tcga_pan_can_atlas_2018); CGGA (http://www.cgga.org.cn). Analysis scripts are available from the corresponding author upon reasonable request.

## Authors’ contributions

YB (Yang Bai) contributed to single-cell RNA-seq data acquisition, expression analysis pipeline development, and figure preparation. HX (Huan Xia) contributed to survival analysis pipeline implementation, Cox regression methodology, and meta-analysis. FW (Fan Wu) provided guidance on AI technology. XT (Xiang Tan) contributed to clinical annotation curation, cohort data harmonization, and manuscript revision. XMW (Xinmin Wu) conceived and designed the study, supervised all analyses, interpreted the results, and wrote the manuscript. All authors read and approved the final manuscript.

## Acknowledgements

The authors thank the Neftel laboratory for making scRNA-seq data publicly available, the TCGA Research Network and CGGA consortium for open-access clinical and transcriptomic datasets, and acknowledge the use of large language models for code debugging and language polishing during manuscript preparation.

## Supplementary Information (Includes Supplementary Figures S1–S6)

**Supplementary Figure S1.**
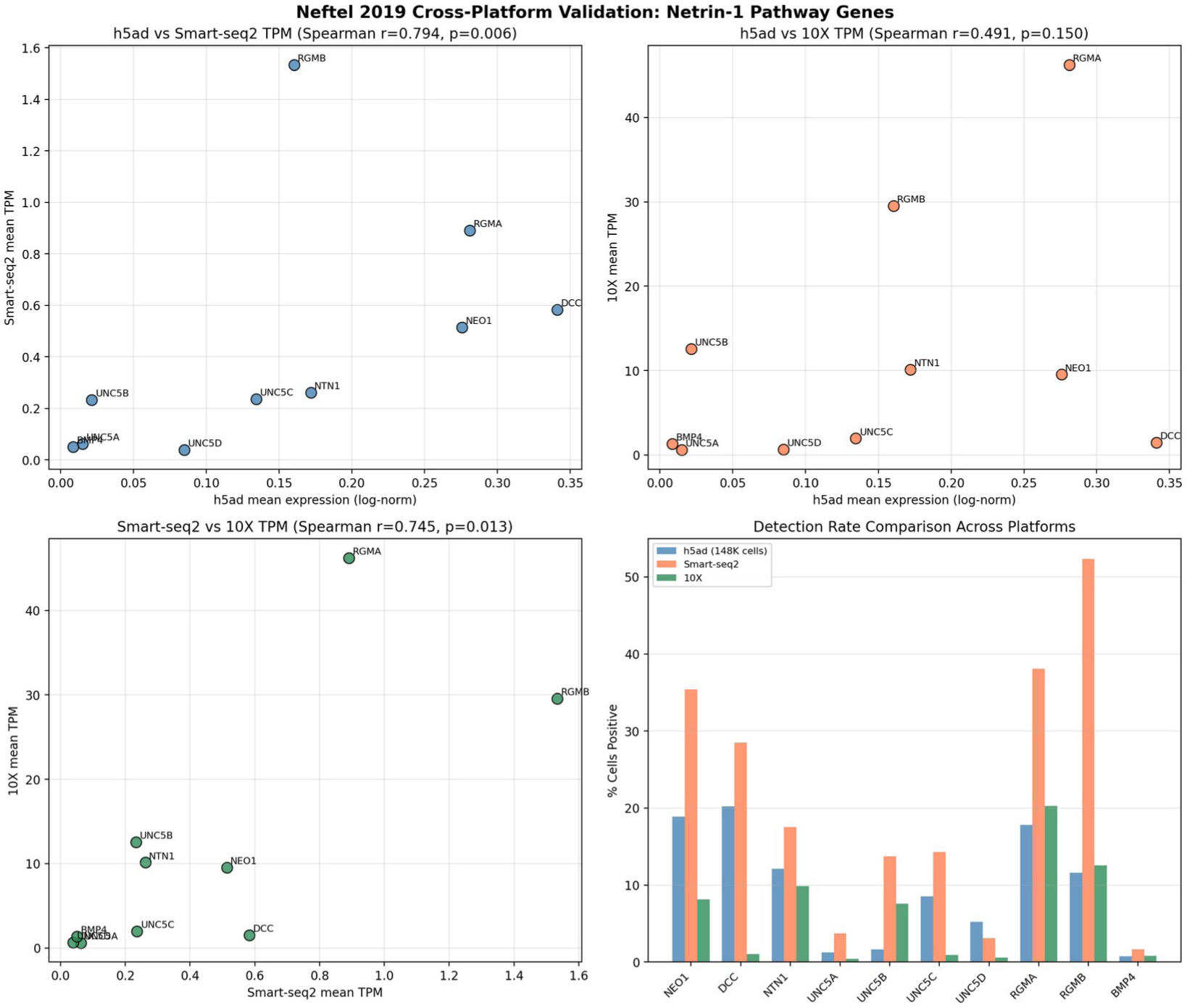
Cross-platform validation of Netrin-1 pathway gene expression. Scatter plots comparing mean expression of 10 pathway genes across three platforms: (top-left) h5ad (SMART-Seq2, 148,019 cells) vs Smart-seq2 single-tumor (7,930 cells), (top-right) h5ad vs 10X Genomics single-tumor (16,201 cells), (bottom-left) Smart-seq2 vs 10X Genomics. Spearman and Pearson correlation coefficients shown. (bottom-right) Detection summary: percentage of 10 pathway genes detected (TPM>1) in each platform.

**Supplementary Figure S2.**
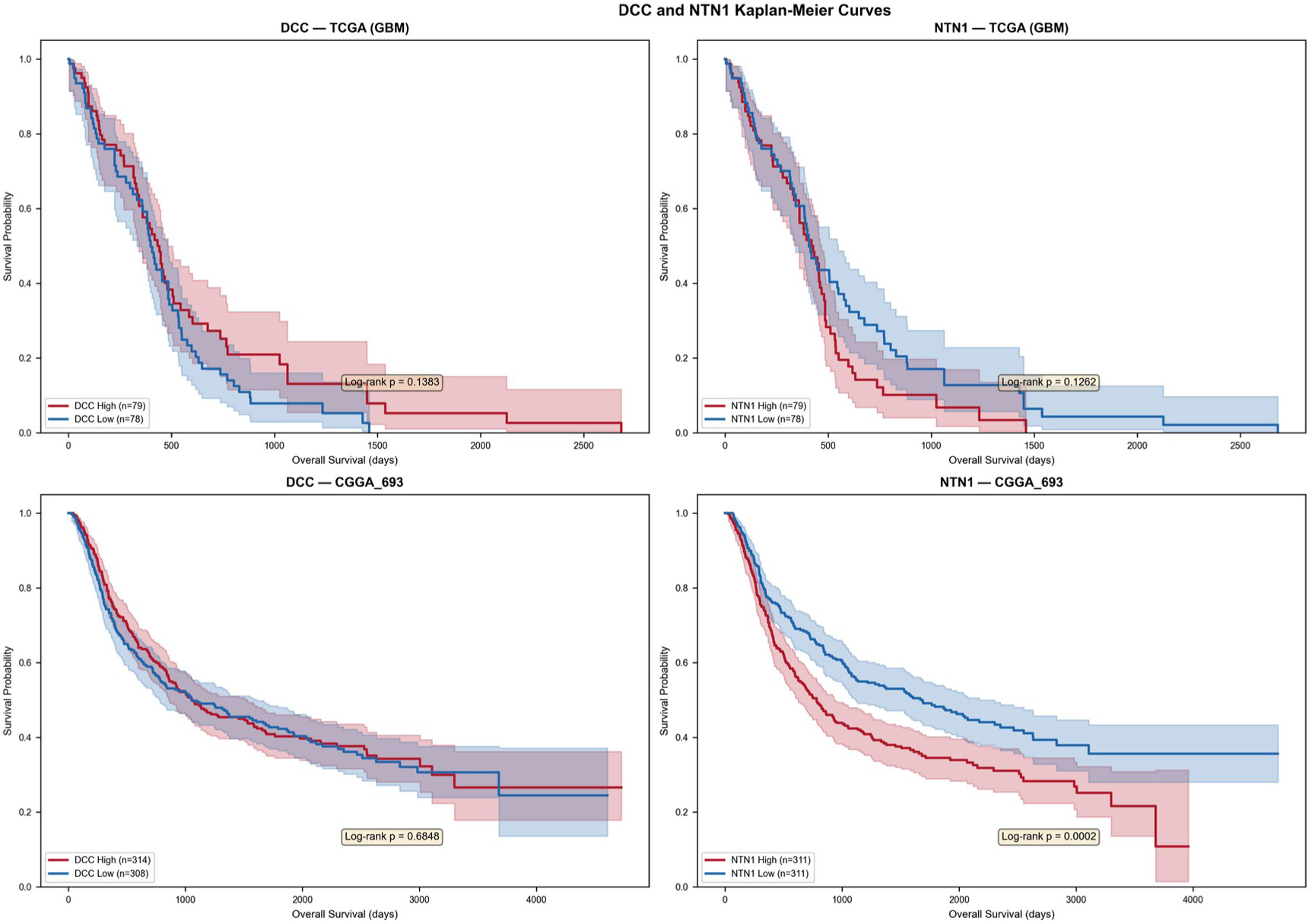
Kaplan-Meier survival curves for Netrin-1 pathway genes in TCGA and CGGA_325 cohorts. Patients stratified by quartile split (Q4=High vs Q1=Low) for NEO1, NTN1, DCC, and RGMB. Log-rank p-values shown for each comparison. Curves demonstrate consistent directional trends across independent cohorts.

**Supplementary Figure S3.**
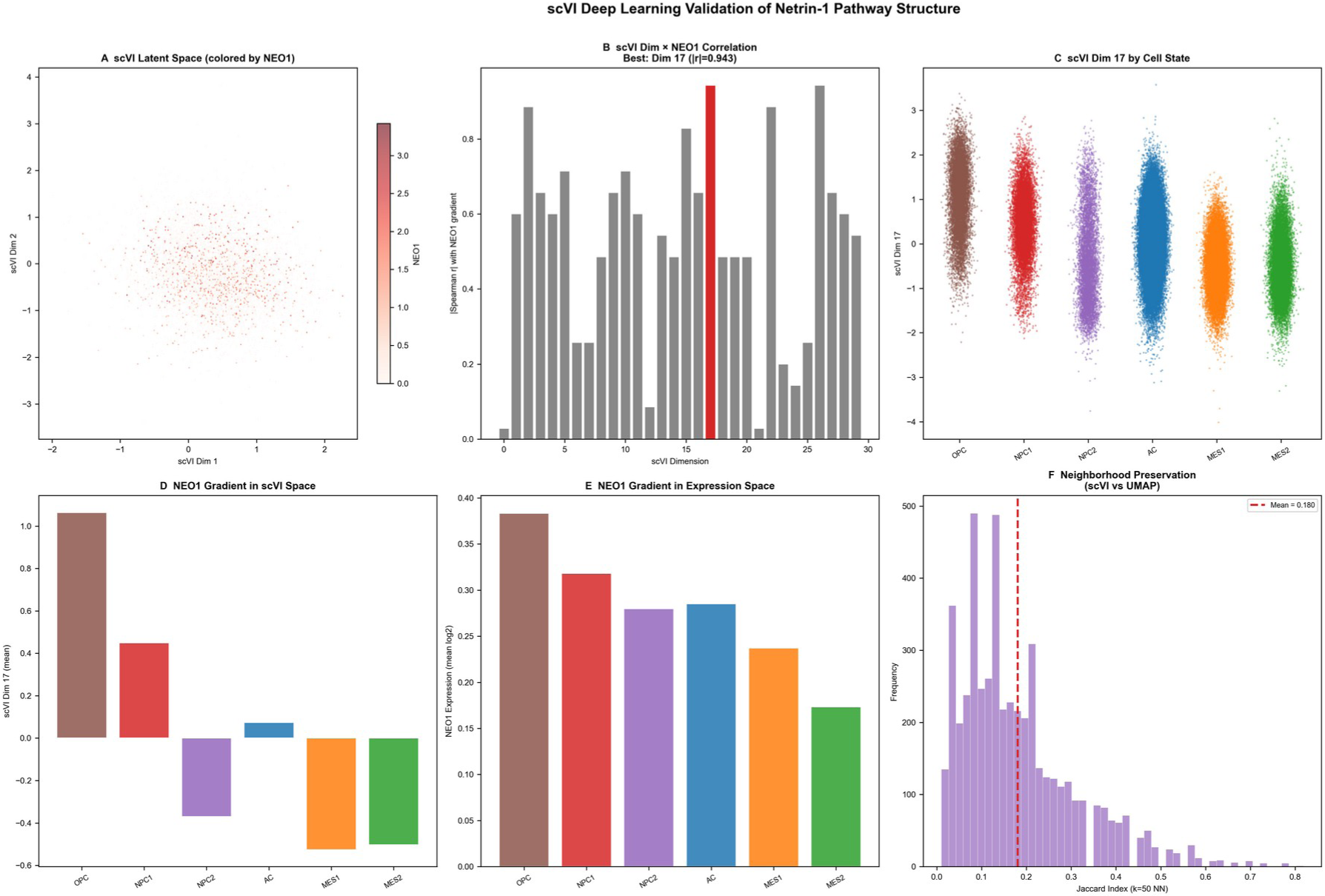
scVI deep learning validation of Netrin-1 pathway structure. (A) scVI latent space (dimensions 1-2) colored by NEO1 expression, demonstrating gradient preservation. (B) Spearman correlation between each scVI dimension and the NEO1 expression gradient across six cellular states. Dimension 17 shows the strongest correlation (|r| =0.943, red bar). (C) scVI dimension 17 values by cell state, mirroring the OPC-to-MES NEO1 gradient. (D) Mean NEO1 gradient representation in scVI space. (E) Mean NEO1 expression gradient in original expression space for comparison. (F) Jaccard neighborhood overlap (k=50 nearest neighbors) between scVI and UMAP embeddings; mean Jaccard index = 0.180 indicates substantial preservation of local transcriptomic structure.

**Supplementary Figure S4.**
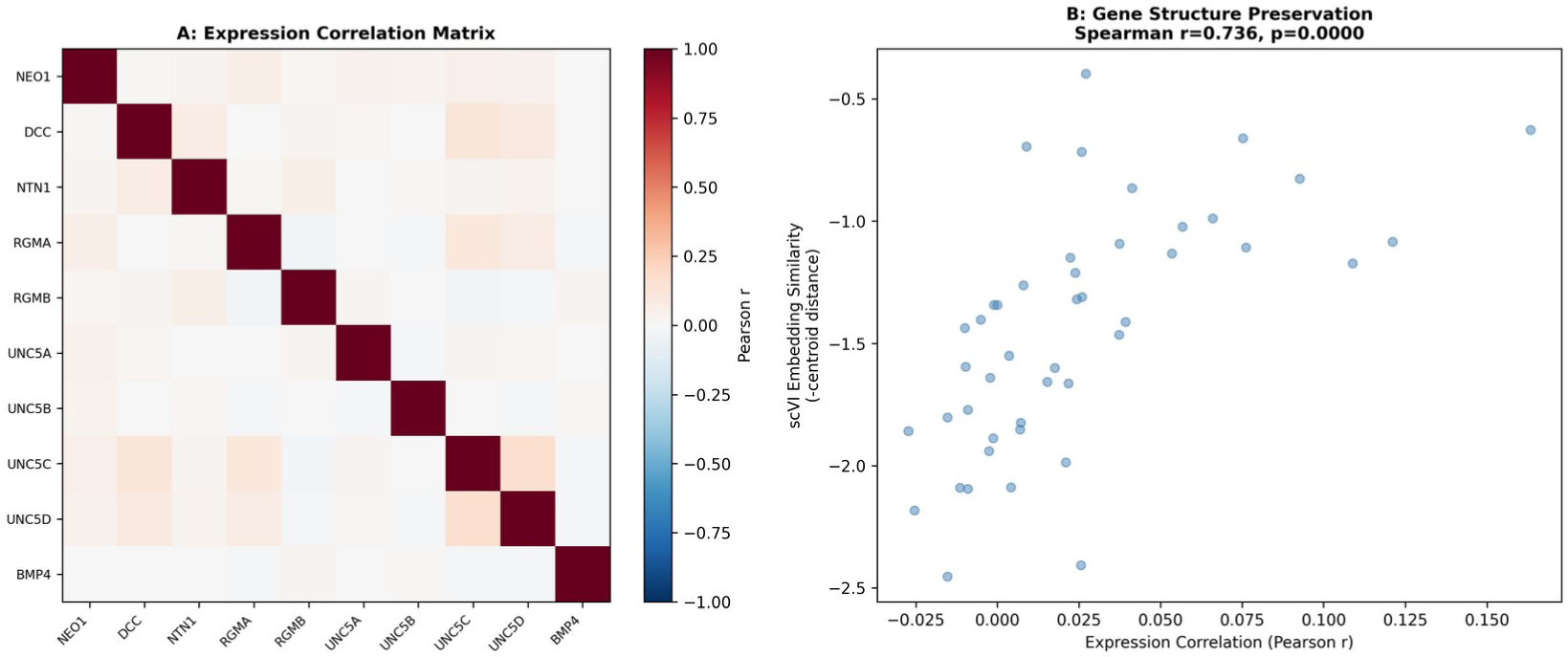
scVI validation of Netrin-1 pathway gene-gene structure preservation. Comparison of pairwise Pearson correlation matrices for 10 Netrin-1 pathway genes in expression space (upper triangle) and scVI latent space (lower triangle). Spearman rank correlation between the two correlation profiles (r=0.736, p<0.0001) confirms that the pathway gene-gene correlation architecture is robust to the choice of dimensionality reduction method.

**Supplementary Figure S5.**
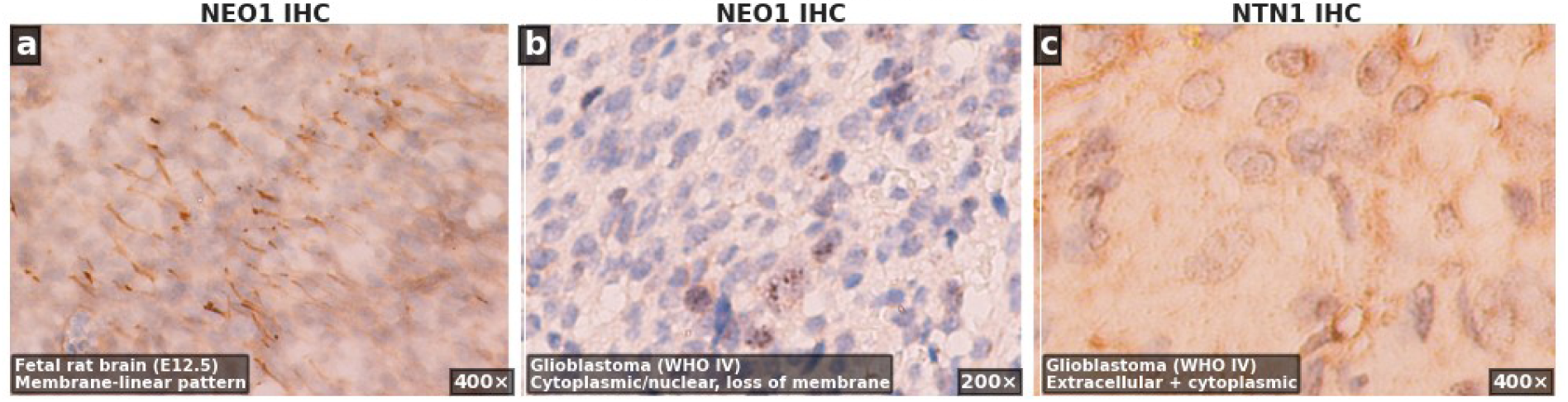
Immunohistochemical localization of NEO1 and NTN1 proteins. (a) NEO1 immunostaining in fetal rat brain (E12.5, 400×) showing a characteristic linear membrane-bound pattern, consistent with its canonical topology as a type I transmembrane dependence receptor. (b) NEO1 immunostaining in human glioblastoma (WHO grade IV, 200×) demonstrating predominantly cytoplasmic and nuclear immunoreactivity with loss of the membrane-linear pattern, suggesting aberrant subcellular trafficking or retention. (c) NTN1 immunostaining in human glioblastoma (WHO grade IV, 400×) revealing extracellular and cytoplasmic distribution, consistent with its identity as a secreted ligand. IHC protocols and antibody details were described previously [13].

**Supplementary Figure S6.**
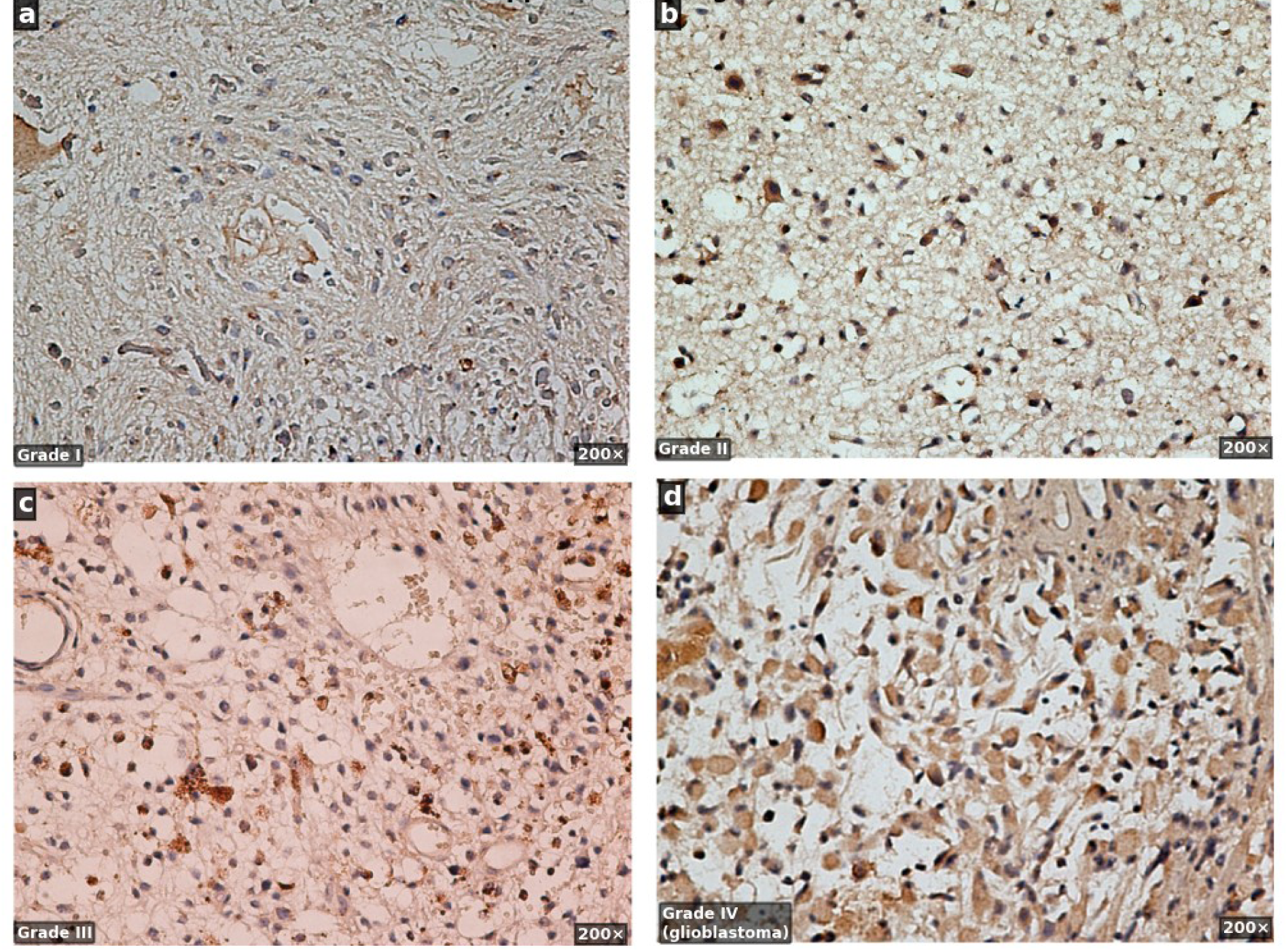
NTN1 immunohistochemical expression across glioma grades. Representative NTN1 immunostaining in (a) WHO grade I, (b) WHO grade II, (c) WHO grade III, and (d) WHO grade IV (glioblastoma) glioma tissues. All panels at 200× original magnification. NTN1 immunoreactivity is detectable across all glioma grades, with increasing extracellular and cytoplasmic distribution associated with higher tumor grade. IHC protocols and antibody details were described previously [13].

**Supplementary Table 1.**
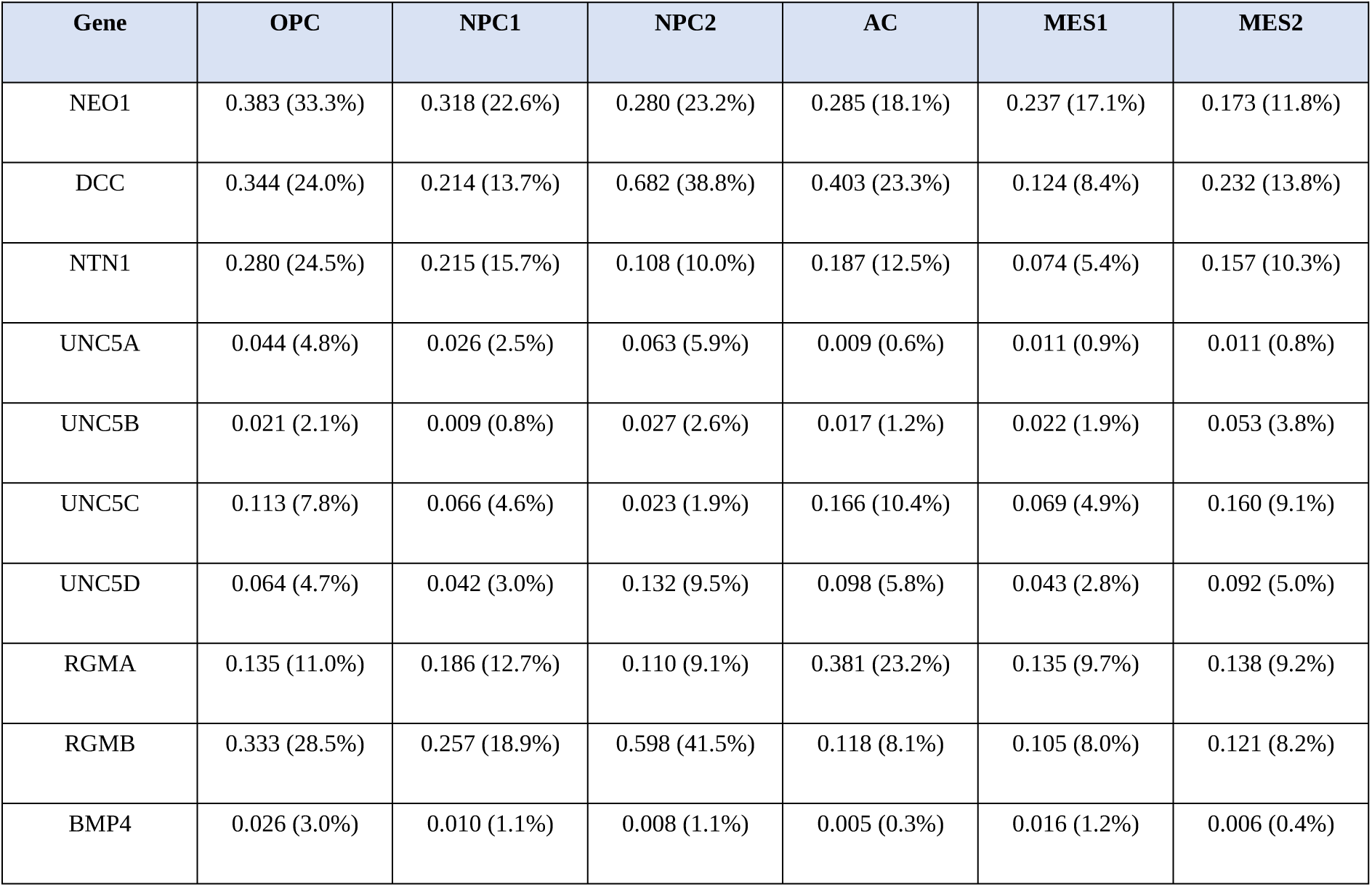
Expression of Netrin-1 pathway genes across GBM cellular states. Mean log-normalized expression and percentage of positive cells for 10 pathway genes across six Neftel cellular states.

**Supplementary Table 2.** Complete subgroup-stratified Cox regression results. Per-SD HR with 95% CI for NEO1, DCC, NTN1, RGMB, and UNC5B in the CGGA_693 cohort.

| Subgroup | Gene | N | Events | HR | 95% CI | p-value |
| --- | --- | --- | --- | --- | --- | --- |
| IDH: IDH-Wildtype | NEO1 | 182 | 158 | 1.034 | 0.870–1.230 | 0.7019 |
| IDH: IDH-Wildtype | DCC | 182 | 158 | 0.846 | 0.625–1.145 | 0.2795 |
| IDH: IDH-Wildtype | NTN1 | 182 | 158 | 1.128 | 0.985–1.291 | 0.0810 |
| IDH: IDH-Wildtype | RGMB | 182 | 158 | 1.092 | 0.914–1.305 | 0.3330 |
| IDH: IDH-Wildtype | UNC5B | 182 | 158 | 1.032 | 0.867–1.227 | 0.7252 |
| IDH: IDH-Mutant | NEO1 | 45 | 32 | 1.093 | 0.720–1.659 | 0.6763 |
| IDH: IDH-Mutant | DCC | 45 | 32 | 0.914 | 0.486–1.717 | 0.7789 |
| IDH: IDH-Mutant | NTN1 | 45 | 32 | 1.367 | 0.979–1.910 | 0.0668 |
| IDH: IDH-Mutant | RGMB | 45 | 32 | 0.837 | 0.571–1.227 | 0.3620 |
| IDH: IDH-Mutant | UNC5B | 45 | 32 | 1.330 | 0.912–1.939 | 0.1382 |
| MGMT: MGMT-Methylated | NEO1 | 104 | 84 | 1.032 | 0.807–1.319 | 0.8019 |
| MGMT: MGMT-Methylated | DCC | 104 | 84 | 0.854 | 0.528–1.379 | 0.5182 |
| MGMT: MGMT-Methylated | NTN1 | 104 | 84 | 1.196 | 1.000–1.430 | 0.0504 |
| MGMT: MGMT-Methylated | RGMB | 104 | 84 | 0.932 | 0.763–1.140 | 0.4961 |
| MGMT: MGMT-Methylated | UNC5B | 104 | 84 | 1.001 | 0.783–1.278 | 0.9952 |
| MGMT: MGMT-Unmethylated | NEO1 | 89 | 78 | 1.125 | 0.861–1.471 | 0.3868 |
| MGMT: MGMT-Unmethylated | DCC | 89 | 78 | 0.924 | 0.643–1.327 | 0.6680 |
| MGMT: MGMT-Unmethylated | NTN1 | 89 | 78 | 1.151 | 0.951–1.394 | 0.1490 |
| MGMT: MGMT-Unmethylated | RGMB | 89 | 78 | 0.986 | 0.776–1.252 | 0.9052 |
| MGMT: MGMT-Unmethylated | UNC5B | 89 | 78 | 1.331 | 1.017–1.741 | 0.0373 |
| Sex: Male | NEO1 | 139 | 117 | 0.917 | 0.753–1.118 | 0.3922 |
| Sex: Male | DCC | 139 | 117 | 1.000 | 0.722–1.383 | 0.9984 |
| Sex: Male | NTN1 | 139 | 117 | 1.136 | 0.974–1.325 | 0.1043 |
| Sex: Male | RGMB | 139 | 117 | 0.887 | 0.721–1.090 | 0.2531 |
| Sex: Male | UNC5B | 139 | 117 | 1.015 | 0.818–1.258 | 0.8943 |
| Sex: Female | NEO1 | 98 | 80 | 1.417 | 1.074–1.869 | 0.0137 |
| Sex: Female | DCC | 98 | 80 | 0.707 | 0.446–1.123 | 0.1419 |
| Sex: Female | NTN1 | 98 | 80 | 1.249 | 1.037–1.505 | 0.0193 |
| Sex: Female | RGMB | 98 | 80 | 1.065 | 0.874–1.296 | 0.5343 |
| Sex: Female | UNC5B | 98 | 80 | 1.239 | 0.997–1.539 | 0.0532 |
| Age: Age<60 | NEO1 | 173 | 139 | 1.139 | 0.946–1.370 | 0.1686 |
| Age: Age<60 | DCC | 173 | 139 | 0.876 | 0.633–1.211 | 0.4227 |
| Age: Age<60 | NTN1 | 173 | 139 | 1.198 | 1.048–1.370 | 0.0081 |
| Age: Age<60 | RGMB | 173 | 139 | 0.988 | 0.834–1.170 | 0.8879 |
| Age: Age<60 | UNC5B | 173 | 139 | 1.161 | 0.980–1.377 | 0.0847 |
| Age: Age≥60 | NEO1 | 64 | 58 | 0.935 | 0.686–1.275 | 0.6723 |
| Age: Age≥60 | DCC | 64 | 58 | 0.856 | 0.513–1.426 | 0.5494 |
| Age: Age≥60 | NTN1 | 64 | 58 | 1.227 | 0.945–1.594 | 0.1253 |
| Age: Age≥60 | RGMB | 64 | 58 | 0.972 | 0.746–1.267 | 0.8360 |
| Age: Age≥60 | UNC5B | 64 | 58 | 1.105 | 0.785–1.554 | 0.5680 |
| RT: RT-Yes | NEO1 | 193 | 157 | 1.050 | 0.878–1.254 | 0.5945 |
| RT: RT-Yes | DCC | 193 | 157 | 0.867 | 0.646–1.162 | 0.3383 |
| RT: RT-Yes | NTN1 | 193 | 157 | 1.156 | 1.017–1.315 | 0.0269 |
| RT: RT-Yes | RGMB | 193 | 157 | 0.944 | 0.806–1.106 | 0.4743 |
| RT: RT-Yes | UNC5B | 193 | 157 | 1.044 | 0.884–1.233 | 0.6109 |
| RT: RT-No | NEO1 | 32 | 29 | 1.219 | 0.831–1.788 | 0.3113 |
| RT: RT-No | DCC | 32 | 29 | 0.885 | 0.374–2.096 | 0.7814 |
| RT: RT-No | NTN1 | 32 | 29 | 1.345 | 0.911–1.986 | 0.1361 |
| RT: RT-No | RGMB | 32 | 29 | 0.987 | 0.680–1.433 | 0.9452 |
| RT: RT-No | UNC5B | 32 | 29 | 1.659 | 1.096–2.512 | 0.0167 |

**Supplementary Table 3.** Schoenfeld residual test results for proportional hazards assumption. Tests for all 30 gene-cohort continuous Cox regression models (10 genes × 3 cohorts).

| Cohort | Gene | N | Schoenfeld p-value | PH Violated |
| --- | --- | --- | --- | --- |
| TCGA | NEO1 | 154 | 0.0812 | No |
| TCGA | DCC | 154 | 0.7060 | No |
| TCGA | NTN1 | 154 | 0.9144 | No |
| TCGA | UNC5A | 154 | 0.1731 | No |
| TCGA | UNC5B | 154 | 0.8911 | No |
| TCGA | UNC5C | 154 | 0.8228 | No |
| TCGA | UNC5D | 154 | 0.5078 | No |
| TCGA | RGMA | 154 | 0.8661 | No |
| TCGA | RGMB | 154 | 0.5419 | No |
| TCGA | BMP4 | 154 | 0.7441 | No |
| CGGA_325 | NEO1 | 305 | 0.0966 | No |
| CGGA_325 | DCC | 305 | 0.5832 | No |
| CGGA_325 | NTN1 | 305 | 0.3085 | No |
| CGGA_325 | UNC5A | 305 | 0.6588 | No |
| CGGA_325 | UNC5B | 305 | 0.9275 | No |
| CGGA_325 | UNC5C | 305 | 0.1041 | No |
| CGGA_325 | UNC5D | 305 | 0.5348 | No |
| CGGA_325 | RGMA | 305 | 0.0032 | Yes |
| CGGA_325 | RGMB | 305 | 0.0701 | No |
| CGGA_325 | BMP4 | 305 | 0.4270 | No |
| CGGA_693 | NEO1 | 611 | 0.0839 | No |
| CGGA_693 | DCC | 611 | 0.1281 | No |
| CGGA_693 | NTN1 | 611 | 0.8381 | No |
| CGGA_693 | UNC5A | 611 | 0.0280 | Yes |
| CGGA_693 | UNC5B | 611 | 0.3459 | No |
| CGGA_693 | UNC5C | 611 | 0.1799 | No |
| CGGA_693 | UNC5D | 611 | 0.3984 | No |
| CGGA_693 | RGMA | 611 | 0.1709 | No |
| CGGA_693 | RGMB | 611 | 0.0081 | Yes |
| CGGA_693 | BMP4 | 611 | 4.83e-05 | Yes |

